# Unveiling the Evolution of CRISPR Spacer Number: A Phylogenetic Analysis of its Correlation with Repeat Characteristics

**DOI:** 10.1101/2024.05.23.595542

**Authors:** Jia Liu, Rui Huang, Deng-Ke Niu

## Abstract

CRISPR-Cas systems in prokaryotes utilize spacers, segments of DNA acquired from invading phages, to guide immune defense mechanisms. This study investigates the evolution of CRISPR repertoire size by examining its relationships with repeat length, terminal repeat polymorphism, and structural stability in 1,958 bacterial genomes, identifying 5,465 CRISPR arrays. Using CRISPRCasFinder for annotation and RNAfold for predicting RNA secondary structures, we found significant variation in array characteristics. Long-repeat arrays (≥38 bp) showed a significant positive correlation between terminal repeat polymorphism and CRISPR spacer number, a correlation absent in short-repeat arrays (<38 bp), suggesting longer repeats facilitate recombination and spacer loss. Additionally, a negative correlation between repeat length and spacer number across all arrays indicates that longer repeats may accelerate spacer loss. Furthermore, our results show that immune demand significantly influences the evolution of spacer number. Larger CRISPR repertoires correlate with conserved repeat sequences and stable secondary structures, vital for functional arrays under continuous selective pressure. Comparing functional and obsolete CRISPR arrays (orphan arrays in genomes lacking Cas genes) revealed that obsolete arrays have fewer spacers and lower repeat consistency, indicating a degenerative state. By elucidating the factors that shape CRISPR memory size evolution, this research offers strategies to enhance bacterial defenses, mitigate resistance, and improve applications in gene editing and therapeutics.

## Introduction

In the ongoing battle against invading genetic elements such as viruses and plasmids, bacteria have evolved a variety of defense mechanisms (Koonin et al. 2017). Among these, the Clustered Regularly Interspaced Short Palindromic Repeats (CRISPR) and their associated protein-coding genes (CRISPR-Cas) represent a sophisticated adaptive immune system. Characterized by repeats typically ranging from 23 to 47 base pairs with loose dyad symmetry, these elements are interspersed with unique sequences derived from foreign DNA (Pourcel et al. 2020). These sequences are transcribed into precursor RNA molecules, which are then processed into mature CRISPR RNA (crRNA). The crRNA includes a repeat-derived segment that aids complex formation with Cas proteins and a spacer-derived segment crucial for targeting specific genetic invaders (Nussenzweig and Marraffini 2020; Watson et al. 2021). Cas genes, located proximal to the CRISPR array, encode effector proteins that cooperate with crRNA to execute the immune response by identifying and cleaving foreign genetic material, ensuring the specificity and effectiveness of the system. Occasionally, CRISPR arrays are found without associated Cas genes, termed “orphan” arrays (Russel et al. 2020; Shmakov et al. 2020), which may be evolutionary remnants or still play roles in cellular functions beyond conventional immune functions.

Spacers within the CRISPR system serve as molecular memories of past infections, enabling rapid and precise defenses against recurrent or similar threats. The diversity of spacers is crucial as it enhances the system’s ability to recognize and combat a broad spectrum of pathogens. Moreover, a rich spacer repertoire increases the likelihood of incorporating new spacers through “primed adaptation” from DNA fragments resembling existing spacers, thus facilitating rapid adaptation to evolving threats (Gong et al. 2019; Jackson et al. 2019). However, an indiscriminate accumulation of spacers can be detrimental (Martynov et al. 2017; Bradde et al. 2020; Garrett 2021; Zaayman and Wheatley 2022). Excessive old spacers may dilute the concentrations of CRISPR effector complexes armed with newly acquired spacers. One potential mechanism against this excessive buildup is spacer loss via homologous recombination among different repeats within the array, which could significantly reduce the immune repertoire if the terminal repeat participates in recombination (Deveau et al. 2008; Horvath et al. 2008; Gudbergsdottir et al. 2011; Jiang et al. 2013; Rao et al. 2017; Stout et al. 2018; Lam and Ye 2019; Deecker and Ensminger 2020). Terminal repeat polymorphisms largely mitigate this risk of erasing many existing immune memories by differing from the uniform sequences found elsewhere in the array (Garrett 2021).

While terminal repeat polymorphisms reduce the likelihood of catastrophic reductions in the immune repertoire, recombination among repeats located near the ends or middle with those at the front can also critically eliminate essential immune memories. A potential evolutionary strategy to mitigate such losses could involve increasing polymorphisms across all repeats as the immune repertoire expands. Considering this mutational force, the size of immune memories might be positively correlated with polymorphisms across all repeats. Additionally, the length of homologous sequences influences the frequency of recombination, with longer repeats providing more sequence for potential recombination events, thereby possibly increasing their frequency (Fujitani et al. 1995; Hua et al. 1997; Lovett 2004). It is hypothesized that there is a negative correlation between repeat length and immune repertoire size. To explore these hypotheses, we will investigate the relationship between repeat length, homogeneity, and immune repertoire size through a comparative analysis of sequenced bacterial genomes.

Palindromic structures within CRISPR repeat sequences exhibit unique properties that enable the formation of relatively conserved secondary RNA structures. These structures enhance the function and stability of crRNA molecules by facilitating precise base pairing. The stability of RNA secondary structures, quantifiable by minimum free energy (MFE), reflects the thermodynamic favorability of these configurations (Mathews and Turner 2006). A low MFE value indicates a robust secondary structure, which is crucial for the effective functioning of crRNAs within the CRISPR-Cas system. Furthermore, stability grants crRNAs resistance against degradation, prolonging their functional presence in the dynamic cellular environment and maintaining sustained defense against invasive genetic elements such as viruses and plasmids. Using crRNA stability as an indicator of the immune effectiveness of CRISPR-Cas systems, we will assess whether a larger immune repertoire size correlates with a more robust immune response. Additionally, we will examine the effect of immune effectiveness on spacer number evolution by comparing obsolete and functional arrays.

## Materials and Methods

### Bacterial Genome Sequence Data and Phylogenetic Tree

We sourced bacterial genome sequence data from the NCBI Genome Database (https://www.ncbi.nlm.nih.gov/genome/browse#!/prokaryotes/) selecting entries that met the assembly quality standards of complete genomes and chromosomes (accessed April 17, 2023), totaling 40,701 entries. The phylogenetic tree was obtained from the Genome Taxonomy Database (GTDB) (Parks et al. 2022), which utilizes strict quality control measures on prokaryotic genome data to construct a comprehensive prokaryotic phylogenetic tree. From GTDB, we utilized the bacterial phylogenetic tree corresponding to the bac120.tree file, containing 62,291 bacterial genome entries (https://data.gtdb.ecogenomic.org/releases/latest/bac120.tree). Cross-referencing the NCBI database, we identified 5,588 entries with matching genome accession numbers, from which 1,958 genomes containing level 4 CRISPR arrays were identified through CRISPRCasdb (Pourcel et al. 2020).

### Annotation of the CRISPR-Cas systems

Using CRISPRCasFinder (Couvin et al. 2018), we annotated CRISPR arrays and associated Cas genes in the 1,958 genomes. This tool provides detailed information on each array, including a consensus sequence of the repeats, which is the highly conserved sequence repeated within the array. This consensus sequence is crucial for the proper formation and function of CRISPR RNAs (crRNAs) and the overall functionality of the CRISPR-Cas system. Annotations were manually curated by referencing data from CRISPRCasdb (Pourcel et al. 2020) to confirm the reliability of arrays, especially those classified below level 4 but showing similarities to level 4 arrays. This process identified 5,465 reliable arrays. CRISPRDirection from CRISPRCasFinder was used to determine the orientation of these arrays, identifying directional information for 1,901 arrays across 814 genomes. The EBcons (entropy-based conservation) index from CRISPRCasFinder was used to assess repeat sequence consistency within each array.

### Consensus Repeat Sequences’ Secondary Structure Quantification

We utilized RNAfold from the ViennaRNA package (version 2.6.4) (Lorenz et al. 2011) to predict RNA secondary structures of consensus repeat sequences in CRISPR arrays, obtaining corresponding MFE values to infer structural stability.

### Phylogenetic Comparative Methods

We computed Pagel’s λ to evaluate the phylogenetic signal of traits using phylosig in the phytools package (Revell, 2012). The results indicated that most traits were significantly influenced by phylogenetic relationships (Supplementary Table 1). To account for phylogenetic autocorrelation, all analyses in this study employed phylogenetic comparative methods. PGLS regression analyses were performed using various evolutionary models in the phylolm package (Ho and Ane, 2014) to account for phylogenetic relationships, selecting results from the model with the lowest AIC value for presentation.

According to the research by Chen et al. (2023), swapping the dependent and independent variables in PGLS regression analysis may lead to different results. Therefore, in all PGLS regression analyses, except for cases where discrete variables must be used as independent variables, we performed analyses with both configurations. Although the slope and *p*-value might change, the direction (positive or negative) and the significance of the results remained consistent. All PGLS regression results consistently indicated significant positive correlation, significant negative correlation, or no correlation. Therefore, one configuration of each pair of results was selected for presentation and subsequent discussion.

### Random Sampling

When performing the same analysis on different samples, different correlation results might be observed due to inherent differences between the two samples or the random noise in the small sample. In this study, we used the sample function in R (version 4.2.1) to randomly draw samples without replacement from the large dataset, matching the number of data points in the smaller dataset. Subsequently, multiple PGLS regression analyses were conducted to verify whether the sample size affects the study results.

## Results

### Overview of CRISPR Array Characteristics

Our analysis of 1,958 bacterial genomes revealed a significant variation in CRISPR array characteristics, with a total of 5,465 arrays identified (Table 1). Most arrays exhibited highly conserved repeat sequences, with 93.8% associated with Cas protein genes. The repeat lengths ranged from 23 to 50 base pairs, roughly following a normal distribution.

**Table 1.** Descriptive statistics of the CRISPR characteristics analyzed in this study.

| Characters | *n* | Median | Mean | Standard deviation | Maximum | Minimum |
| --- | --- | --- | --- | --- | --- | --- |
| Array number | 1,958 | 2 | 2.79 | 2.77 | 28 | 1 |
| Spacer number | 5,465 | 16 | 27.1 | 34.4 | 587 | 1 |
| Repeat length | 5,465 | 32 | 32.7 | 4.19 | 50 | 23 |
| EBcons | 5,465 | 96.7 | 93.9 | 7.88 | 100 | 33.8 |

### Effect of Recombination: Evidence from Terminal Repeat Polymorphism

It is generally believed that the polymorphism of terminal repeat sequences is significant because it can effectively prevent the loss of spacer sequences due to homologous recombination. From this, it can be inferred that lower terminal repeat polymorphism may lead to higher recombination efficiency with other repeat sequences within the CRISPR array, potentially resulting in more spacer sequence loss.

In this study, we used the number of base differences between the terminal repeat sequences and the consensus repeat sequence as an indicator of terminal repeat polymorphism. The more base differences in the terminal repeat sequence, the higher the polymorphism. In the 814 genomes with identified CRISPR array orientations, we divided them into low polymorphism and high polymorphism groups based on the number of nucleotide differences between the terminal repeat and the consensus sequence. Nine thresholds, from 0 to 8, were used for categorization (Table 2). In cases where multiple arrays were present in the same genome, we used the median value of the nucleotide differences between the terminal repeat and the consensus sequence to represent the terminal repeat polymorphism of the genome.

**Table 2.** PGLS Results of the Relationship Between Terminal Repeat Polymorphism and Spacer Number

| Polymorphism Threshold | Low Polymorphism | High Polymorphism | Slope | *p* |
| --- | --- | --- | --- | --- |
| 0 | 283 | 531 | 1.445 | 0.480 |
| 1 | 397 | 417 | 1.305 | 0.471 |
| 2 | 481 | 333 | 0.621 | 0.714 |
| 3 | 554 | 260 | 0.962 | 0.566 |
| 4 | 628 | 186 | 2.141 | 0.265 |
| 5 | 679 | 135 | 2.362 | 0.250 |
| 6 | 726 | 88 | 3.248 | 0.192 |
| 7 | 751 | 63 | -0.471 | 0.865 |
| 8 | 776 | 38 | -2.614 | 0.454 |

We then analyzed the correlation between spacer number and terminal repeat polymorphism in multiple groups using PGLS regression analysis. A positive and significant slope indicates a significant positive correlation between the variables, while a negative and significant slope indicates a significant negative correlation. If the slope’s p-value is not below the significance level (usually 0.05), there is no statistically significant correlation between the variables. We found that the spacer number in CRISPR arrays is not significantly correlated with terminal repeat polymorphism (p-values are all greater than 0.1, Table 2).

We hypothesize that homologous recombination is more likely to occur in CRISPR arrays with longer repeat sequences, leading to the loss of spacer sequences. Most repeats may be too short to serve as effective substrates for homologous recombination. To test this possibility, we divided the 814 bacterial genomes into a long-repeat subsets (≥38 bp) and the short-repeat subset (<38 bp). This choice aims to make repeat sequences of the long-repeat subsets as long as possible while ensuring sufficient sample size. If a genome contains both long-repeat arrays and short-repeat arrays, its arrays will be separated into the long-repeat subset and the short-repeat subset. Therefore, the total sample size of the two subset is larger than 814.

Within both the long-repeat subset and the short-repeat subset, we repeated the analysis producing Table 2. In the ≥38 bp group, we could see a significant positive correlation between terminal repeat polymorphism and spacer number (*p* < 0.01, Table 3), no matter which threshold was used to separate the low and high polymorphism groups. On the contrary, the short-repeat subset consistently lacks significant correlations (*p* > 0.1, Table 4). These results suggest that homologous recombination may play a role in spacer number evolution in CRISPR arrays with longer repeat sequences.

**Table 3.** PGLS Results of the Relationship Between Terminal Repeat Polymorphism and Spacer Number in Long-Repeat (≥38 bp) Genomes

| Polymorphism Threshold | Low Polymorphism | High Polymorphism | Slope | *p* |
| --- | --- | --- | --- | --- |
| 0 | 7 | 15 | 15.083 | 0.008 |
| 1 | 10 | 12 | 15.696 | 0.007 |
| 2 | 12 | 10 | 16.942 | 0.003 |
| 3 | 14 | 8 | 17.264 | 0.003 |
| 4 | 15 | 7 | 17.810 | 0.001 |
| 6 | 17 | 5 | 17.153 | 0.002 |
| 11 | 18 | 4 | 17.919 | 0.001 |

**Table 4.** PGLS Results of the Relationship Between Terminal Repeat Polymorphism and Spacer Number in Short-Repeat (< 38 bp) Genomes

| Polymorphism Threshold | Low Polymorphism | High Polymorphism | Slope | *p* |
| --- | --- | --- | --- | --- |
| 0 | 280 | 523 | 1.170 | 0.582 |
| 1 | 392 | 411 | 1.024 | 0.584 |
| 2 | 476 | 327 | 0.346 | 0.843 |
| 3 | 546 | 257 | 0.890 | 0.607 |
| 4 | 620 | 183 | 2.109 | 0.289 |
| 5 | 673 | 130 | 2.510 | 0.240 |
| 6 | 720 | 83 | 2.884 | 0.269 |
| 7 | 745 | 58 | -1.078 | 0.710 |
| 8 | 770 | 33 | -3.763 | 0.311 |

We noticed that there are only 22 samples in the long-repeat subset. Random variations in the small dataset possibly lead to the detection of significant correlations that do not actually exist in the population. To exclude this possibility, we performed 1,000 random resamplings of 22 samples from the short-repeat subset. We set the polymorphism threshold to 1, ensuring minimal difference between low and high polymorphism sample sizes. Analyzing the correlations between spacer number and terminal repeat polymorphism, we found that 916 out of 1,000 resamplings showed no significant correlation, excluding sample size effects in the results shown in Table 3.

Besides 38 bp, we also used 40 bp and 36 bp as thresholds for subsequent calculations. With 40 bp, results were consistent with 38 bp but with a smaller long-repeat sample size (13 bacterial genomes, Supplementary Tables 5-6). With 36 bp, results were not significant, possibly due to the presence of shorter repeat sequences less prone to recombination (Supplementary Tables 7-8).

### Effect of Recombination: Evidence from Repeat Length

The long-repeat subset and the short-repeat subset exhibit a significant difference in the correlation between the number of spacers and terminal repeat polymorphism, suggesting that longer repeat lengths may accelerate spacer loss by increasing recombination efficiency. To further test this idea, we used PGLS regression to examine the relationship between repeat length and the number of spacers in a larger dataset (not distinguishing array orientation, 1958 bacterial genomes). We found a significant negative correlation (*p* = 3.0×10^−10^, Figure 1). The longer the CRISPR repeat length, the fewer the spacer sequences.

**Figure 1.**
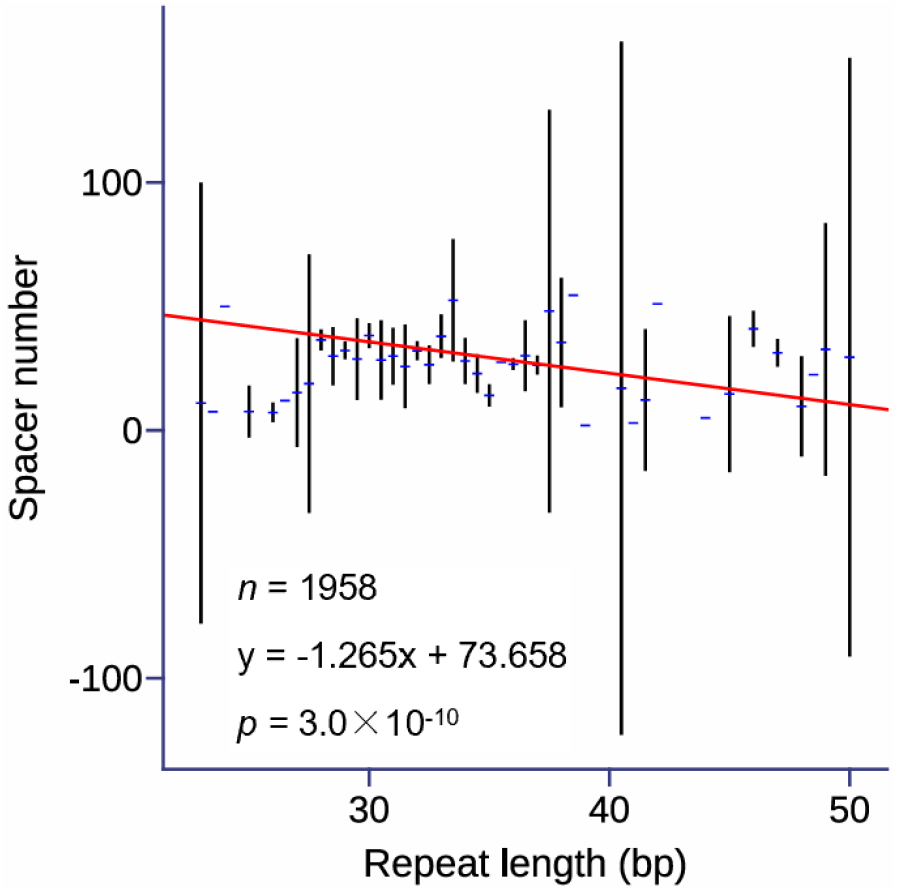
Relationship between CRISPR repeat length and spacer number. The red line represents the regression line obtained from the PGLS regression. Due to phylogenetic correction, the regression line may not visually align with the overall trend perceived intuitively. Error bars indicate the confidence interval for the mean. The figure displays the results from Pagel’s lambda model, which had the lowest AIC value among the four models tested. Results from other models are presented in Supplementary Table 9.

### Effect of Immune Demand: Evidence from Repeat Characteristics

In addition to the possibility of terminal repeat sequences undergoing homologous recombination with internal sequences within CRISPR arrays, leading to the loss of spacer sequences, homologous recombination among other internal repeat sequences within the array may also influence the loss of spacers. Therefore, this study employed Ebcons to measure the consistency among all repeats within the CRISPR arrays. It is hypothesized that higher Ebcons values indicate higher recombination efficiency among repeats within the array, potentially leading to more frequent loss of spacer sequences. However, PGLS analysis of the relationship between the number of spacer sequences and Ebcons revealed a significant positive, rather than negative, correlation between the quantity of CRISPR spacer sequences and the consistency of CRISPR repeat sequences (*p* = 4.8×10^−32^, Figure 2A).

**Figure 2.**
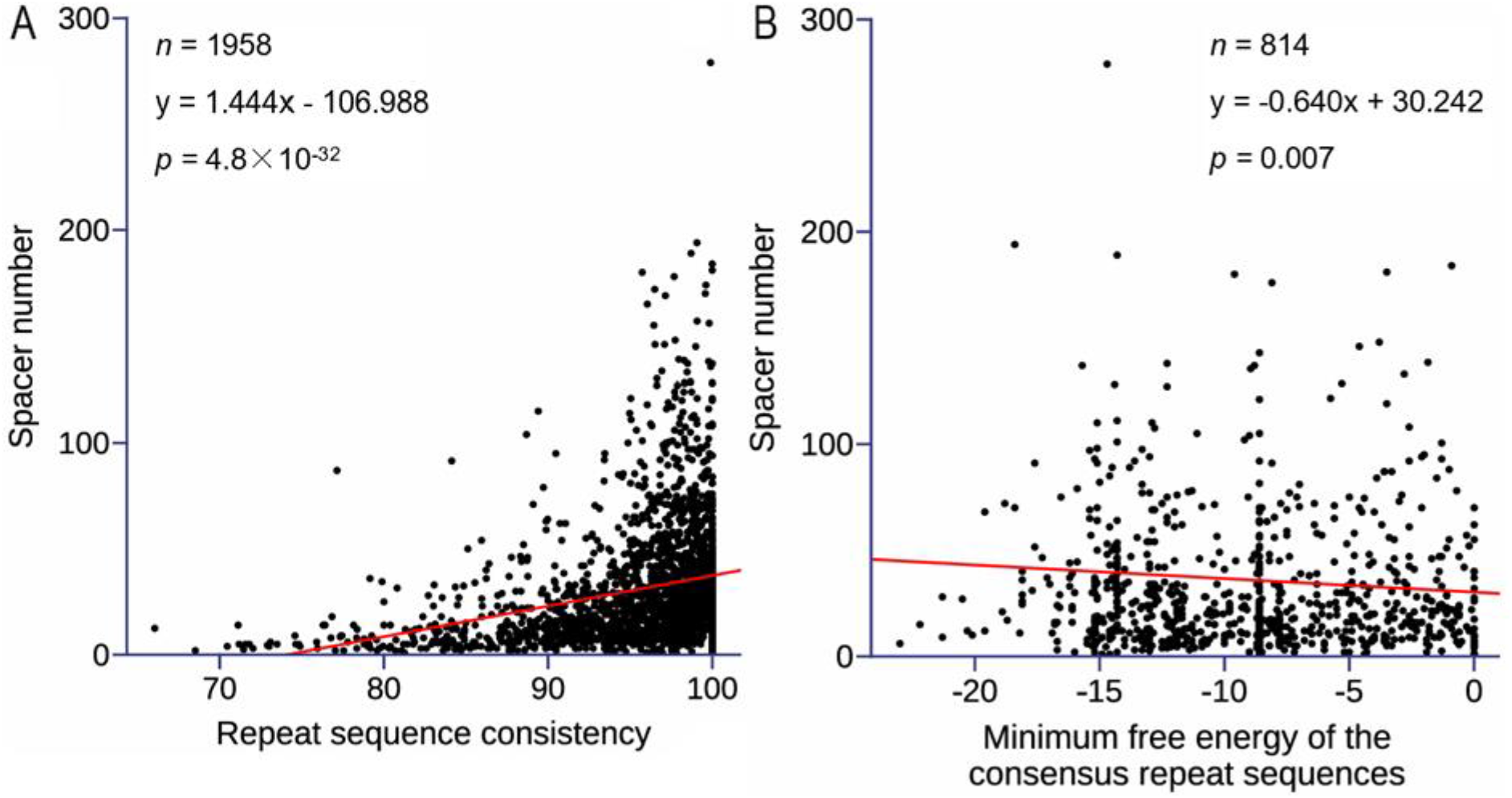
Relationship between spacer number and repeat sequence characteristics. (A) Repeat sequence consistency as represented by the Ebcons (entropy-based conservation). The figure shows the results using Pagel’s lambda model, which had the lowest AIC value among the four models tested. Results from other models are presented in Supplementary Table 10. (B) Secondary structure stability of consensus repeat sequences as represented by the minimum free energy of the consensus sequences. The figure shows the results using the OU model, which has the lowest AIC value among the four models we used. Other model results are in Supplementary Table 11. The red line represents the regression line obtained from the PGLS regression. Due to phylogenetic correction, the regression line may not visually align with the overall trend perceived intuitively.

Contrary to initial expectations, we hypothesize that some CRISPR arrays may be in a process of evolutionary degeneration, akin to pseudogenization, with a reduced evolutionary pressure against the loss of spacer sequences. Changes within the CRISPR repeat sequences could hinder the correct folding of crRNA or its effective binding with Cas proteins, thereby reducing the system’s capability to recognize and cut foreign DNA. On the other hand, bacteria that continuously face repeated attacks from various genetic invaders need to transcribe crRNA and form complexes with Cas proteins for an immune response, thus requiring conserved repeat sequences and numerous spacers.

The more stable the secondary structure of CRISPR repeat sequences, the more stable the crRNAs formed, and the more effectively they can perform their immune functions. Therefore, bacteria that continually face multiple genetic invaders need not only numerous spacer sequences but also repeat sequences with stable secondary structures. We regard the minimum free energy (MFE) of the consensus repeat sequence as an indicator of the stability of the repeat’s secondary structure, where a lower MFE value typically indicates a more stable structure because it requires less energy to form. PGLS regression analysis found a significant negative correlation between the consensus repeat sequence MFE and the number of CRISPR spacer sequences (*p* = 0.007, Figure 2B), reflecting that the stable secondary structure of consensus repeat sequences might be driven by similar selective pressures as the number of spacer sequences.

### Changes in Spacer and Repeat Characteristics during CRISPR Arrays Degeneration

Following the above rationale, obsolete CRISPR arrays, which likely have lost their adaptive immune functions, should show lower numbers of spacer sequences and less functionally constrainted repeat sequences compared to functional arrays. Despite being devoid of nearby Cas genes, orphan arrays may still tap into Cas machinery generated by distant genes or even Cas genes residing on separate DNA molecules (Deng et al. 2013; Bernheim et al. 2019; Deecker and Ensminger 2020; Shmakov et al. 2020; Pinilla-Redondo et al. 2021). Consequently, we classify obsolete CRISPR arrays as orphan arrays found in genomes lacking any Cas genes. These arrays likely represent vestiges of ancestral CRISPR systems, having forfeited their adaptive immune functions over evolutionary time. This stringent classification enables a definitive demarcation between functional and non-functional arrays, thereby facilitating robust comparative analyses.

We find that obsolete CRISPR arrays harbor significantly fewer spacer sequences and display lower consistency among CRISPR repeat sequences compared to functional arrays (*p* = 2.1×10^−9^ and p = 8.9×10^−15^, respectively, as shown in Figure 3A-B). Moreover, examination of secondary structures unveils that the consensus repeat sequences in obsolete arrays are less stable, albeit not statistically significant, compared to functional counterparts (Figure 3C).

**Figure 3.**
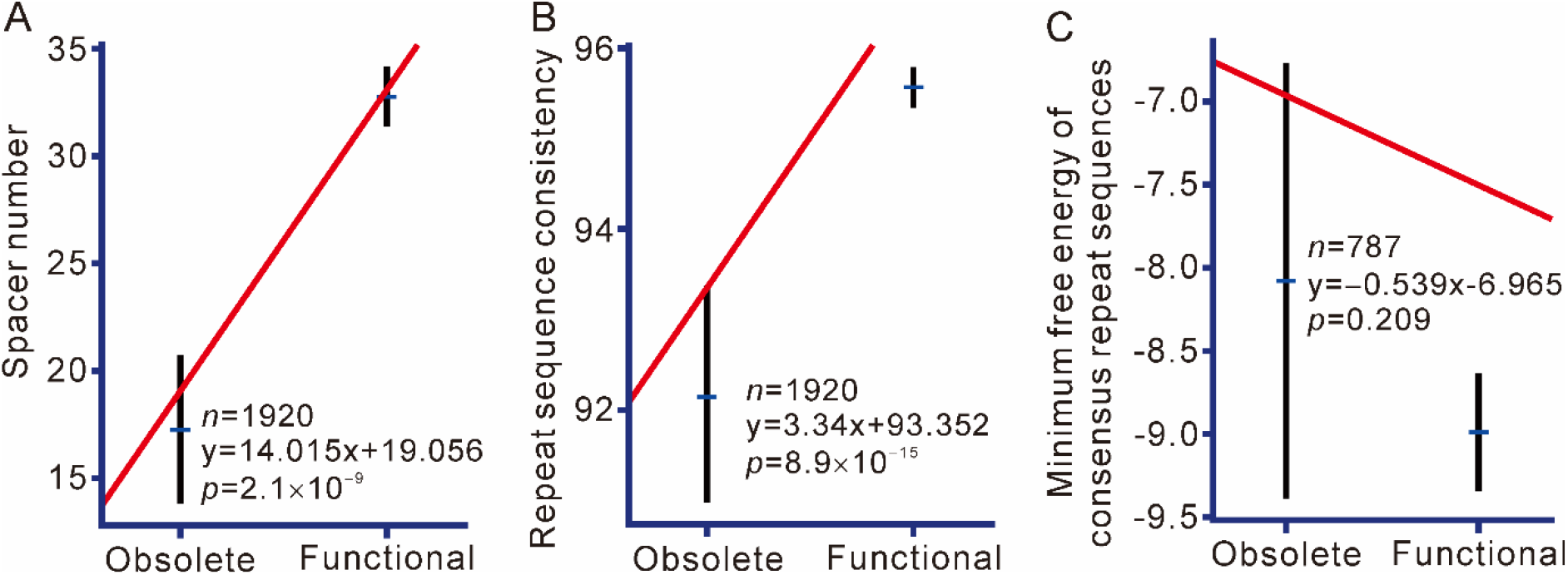
Comparing obsolete arrays with functional arrays. (A) Spacer number comparison between obsolete and functional arrays. (B) Repeat sequence consistency comparison between obsolete and functional arrays, represented by Ebcons (entropy-based conservation). (C) Comparison of consensus repeat sequence secondary structure stability between obsolete and functional arrays, represented by the minimum free energy of the consensus sequences. The red line represents the regression line obtained from the PGLS regression. Due to phylogenetic correction, the regression line may not visually align with the overall trend perceived intuitively. Error bars indicate the confidence interval for the mean. The figure shows the results using Pagel’s lambda model, which had the lowest AIC value among the four models tested. Results from other models are presented in Supplementary Tables 12-13.

Furthermore, our investigation into the relationships between spacer number, repeat sequence consistency, and secondary structure stability uncovers intriguing patterns. While both obsolete and functional arrays exhibit a positive correlation between spacer number and repeat sequence consistency, the relationship is notably weaker in obsolete arrays, suggesting a diminished ability to maintain array integrity (Figure 4A-B). More strikingly, whereas functional arrays demonstrate a positive correlation between spacer number and secondary structure stability, obsolete arrays lack a significant negative correlation, hinting at potential structural destabilization in these evolutionary relics (Figure 4C-D). This pattern not only underscores the degenerative state of these arrays but also serves as indirect evidence of how immune demand influences the evolutionary trajectory of spacer numbers.

**Figure 4.**
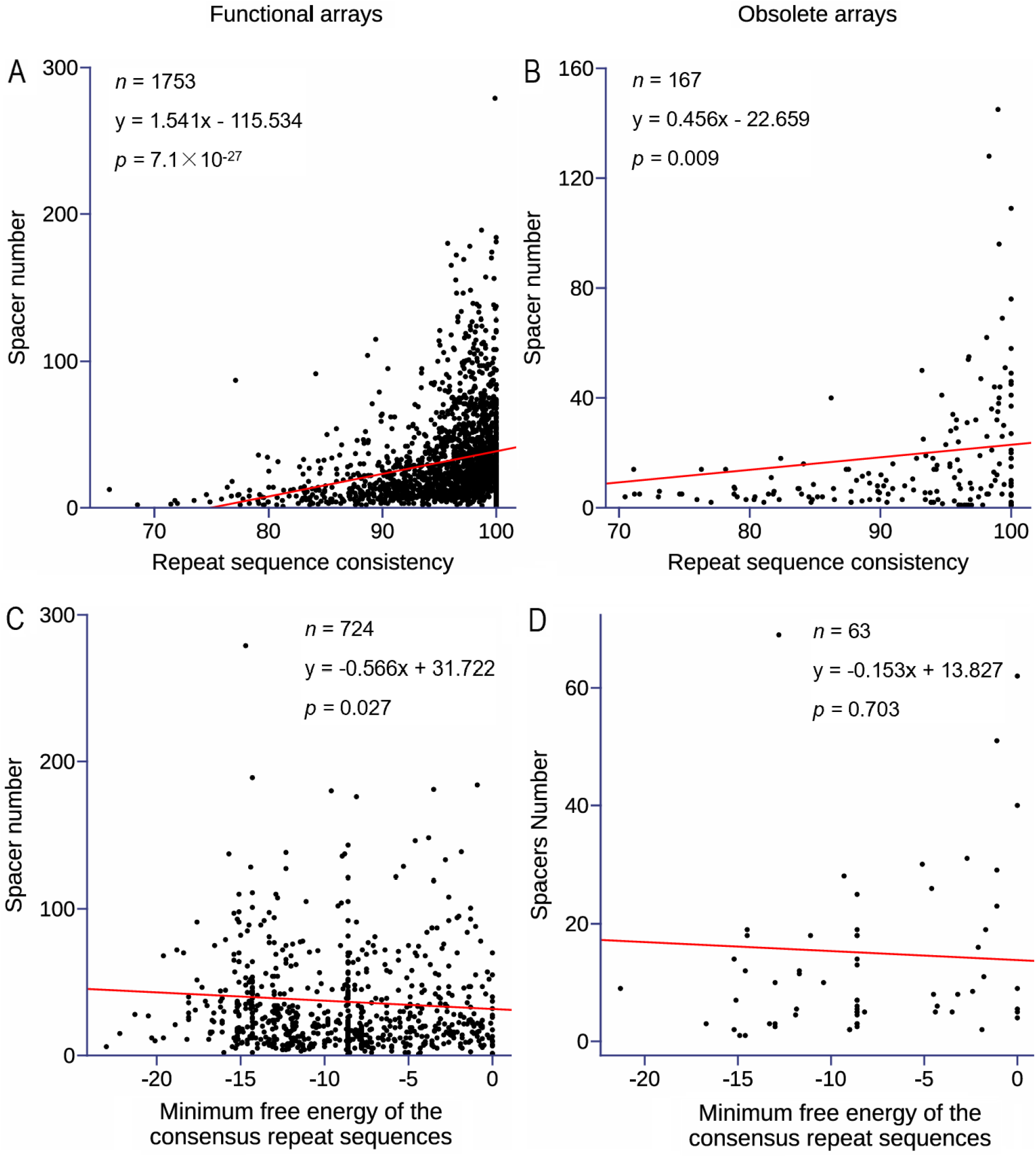
Relationship between spacer number and repeat characteristics. (A) Repeat sequence consistency as measured by Ebcons (entropy-based conservation) in functional arrays. (B) Repeat sequence consistency as measured by Ebcons in obsolete arrays. (C) Stability of the secondary structure of consensus repeat sequences as measured by the minimum free energy of the consensus sequences in functional arrays. (D) Stability of the secondary structure of consensus repeat sequences as measured by the minimum free energy of the consensus sequences in obsolete arrays. The red line represents the regression line obtained from the PGLS regression. Due to phylogenetic correction, the regression line may not visually align with the overall trend perceived intuitively. Error bars indicate the confidence interval for the mean. (A) and (B) show the results using OU model while (C) and (D) show the results using Pagel’s lambda model. These results are presented here because of their lowest AIC value among the four models we used. Other model results are in Supplementary Table 14-15.

## Discussion

This study explores the evolutionary dynamics of CRISPR arrays, focusing on the relationships between spacer number, repeat length, terminal repeat polymorphism, and structural stability. We analyzed 1958 bacterial genomes, identifying 5465 CRISPR arrays through detailed annotation using CRISPRCasFinder, and revealed significant variation in CRISPR array characteristics. While no significant correlation was found between spacer number and terminal repeat polymorphism in the general dataset, a significant positive correlation emerged in CRISPR arrays with longer repeats (≥38 bp), indicating that longer repeats may facilitate recombination and spacer loss. Conversely, the short-repeat subset (<38 bp) showed no significant correlations. Additionally, a significant negative correlation was observed between repeat length and spacer number, suggesting that longer repeats might accelerate spacer loss through recombination.

Our findings also indicate that immune demand is a significant factor influencing spacer number evolution. A higher spacer number is associated with conserved repeat sequences and stable secondary structures of the consensus repeat sequences. Functional arrays, which are subject to continuous selective pressure from genetic invaders, display positive correlations between spacer number and both repeat sequence consistency and secondary structure stability. In contrast, obsolete arrays show weaker correlations for repeat sequence consistency and no significant correlation for structural stability, highlighting their degenerative state and reduced evolutionary pressure.

Previous theoretical studies have suggested that the range of 10-40 spacers within CRISPR arrays provides an optimal defense against a broad spectrum of phages under various realistic parameter estimates (Martynov et al. 2017; Bradde et al. 2020). This aligns with the idea that spacer numbers are balanced by multiple evolutionary forces, such as recombination and immune demand. Another force shaping the evolution of spacer numbers in CRISPR arrays is the need to avoid autoimmunity, as proposed by Chen et al. (2022). Their study revealed a significant positive correlation between spacer length and CRISPR spacer number, suggesting that the risk of autoimmunity resulting from a large number of spacers is mitigated by longer spacers, which reduce the possibility of matching bacterial genome sequences.

These insights enhance our understanding of CRISPR array evolution and the factors influencing spacer retention and loss. The research provides valuable knowledge for improving CRISPR technology and developing strategies to bolster bacterial defenses or counteract bacterial resistance, with potential implications for gene editing and therapeutic applications.

## Notes

### Competing Interest Statement

The authors have declared no competing interest.

### Summary of Updates

The manuscript title has been improved to more accurately reflect the contents.

