## Supplemental Tables for "Unveiling the Evolution of CRISPR Spacer Number: A Phylogenetic Analysis of its Correlation with Repeat Characteristics"

Supplementary Table 1. phylogenetic signal of traits

| *n* | Trait | Pagel’s λ | *p* |
| --- | --- | --- | --- |
| 1,958 | Spacer number | 0.683 | 4.2×10^-57^ |
| 1,958 | Repeat sequence consistency | 0.264 | 9.8×10^-45^ |
| 1,958 | Repeat length | 0.674 | 7.3×10^-279^ |
| 1,920 | Obsolete arrays with functional arrays | 0.225 | 4.3×10^-32^ |
| 814 | Secondary structure stability of consensus repeat sequences | 0.829 | 1.6×10^-126^ |

The phylogenetic signal of each feature in the largest dataset is displayed.

Supplementary Table 2. PGLS Results of the Relationship Between Terminal Repeat Polymorphism and Spacer Number

| Polymorphism Threshold | Low Polymorphism | High Polymorphism | Model | Slope | *p* |
| --- | --- | --- | --- | --- | --- |
| 0 | 283 | 531 | Lambda | 2.183  1.307  1.307 | 0.308  0.445  0.445 |
|  |  |  | BM |  |  |
|  |  |  | EB |  |  |
| 1 | 397 | 417 | Lambda | 1.081  1.650  1.650 | 0.595  0.251  0.251 |
|  |  |  | BM |  |  |
|  |  |  | EB |  |  |
| 2 | 481 | 333 | Lambda | 1.264  0.186  0.186 | 0.527  0.885  0.885 |
|  |  |  | BM |  |  |
|  |  |  | EB |  |  |
| 3 | 554 | 260 | Lambda | 1.921  0.792  0.792 | 0.346  0.525  0.525 |
|  |  |  | BM |  |  |
|  |  |  | EB |  |  |
| 4 | 628 | 186 | Lambda | 4.833  0.342  0.342 | 0.032  0.813  0.813 |
|  |  |  | BM |  |  |
|  |  |  | EB |  |  |
| 5 | 679 | 135 | Lambda | 4.989  -0.196  -0.196 | 0.051  0.897  0.897 |
|  |  |  | BM |  |  |
|  |  |  | EB |  |  |
| 6 | 726 | 88 | Lambda | 4.719  -0.053  -0.053 | 0.127  0.977  0.977 |
|  |  |  | BM |  |  |
|  |  |  | EB |  |  |
| 7 | 751 | 63 | Lambda | 1.662  -2.690  -2.690 | 0.637  0.179  0.179 |
|  |  |  | BM |  |  |
|  |  |  | EB |  |  |
| 8 | 776 | 38 | Lambda | 0.770 | 0.868 |
|  |  |  | BM | -5.168 | 0.043 |
|  |  |  | EB | -5.168 | 0.043 |

Data includes 814 bacterial genomes together. The threshold for polymorphism is set as the number of base differences between the terminal repeat and the consensus sequence, ranging from 0 to 8. Low polymorphism is defined as less than or equal to the threshold, and high polymorphism is defined as greater than the threshold. We labeled groups with low and no polymorphism as 0 and groups with high polymorphism as 1. In PGLS regression analysis, polymorphism group label is used as the independent variable, and the number of spacer sequences as the dependent variable.

Supplementary Table 3. PGLS Results of the Relationship Between Terminal Repeat Polymorphism and Spacer Number in Long-Repeat (≥ 38 bp) Genomes

| Polymorphism Threshold | Low Polymorphism | High Polymorphism | Model | Slope | *p* |
| --- | --- | --- | --- | --- | --- |
| 0 | 7 | 15 | Lambda | 15.083  15.083  15.438 | 0.008  0.008  0.004 |
|  |  |  | OU |  |  |
|  |  |  | EB |  |  |
| 1 | 10 | 12 | Lambda | 15.696  15.696  15.918 | 0.007  0.007  0.004 |
|  |  |  | OU |  |  |
|  |  |  | EB |  |  |
| 2 | 12 | 10 | Lambda | 17.348  16.942  17.045 | 0.005  0.003  0.002 |
|  |  |  | OU |  |  |
|  |  |  | EB |  |  |
| 3 | 14 | 8 | Lambda | 17.962  17.264  17.288 | 0.005  0.003  0.001 |
|  |  |  | OU |  |  |
|  |  |  | EB |  |  |
| 4 | 15 | 7 | Lambda | 18.886  17.810  17.805 | 0.003  0.001  3.0×10^-4^ |
|  |  |  | OU |  |  |
|  |  |  | EB |  |  |
| 6 | 17 | 5 | Lambda | 17.506  17.153  17.221 | 0.003  0.002  7.7×10^-4^ |
|  |  |  | OU |  |  |
|  |  |  | EB |  |  |
| 11 | 18 | 4 | Lambda | 19.041  17.919  17.875 | 0.003  0.001  3.7×10^-4^ |
|  |  |  | OU |  |  |
|  |  |  | EB |  |  |

Data includes 22 bacterial genomes.The threshold for polymorphism is set as the number of base differences between the terminal repeat and the consensus sequence, ranging from 0 to 4, 6, and 11. Low polymorphism is defined as less than or equal to the threshold, and high polymorphism is defined as greater than the threshold. We labeled groups with low and no polymorphism as 0 and groups with high polymorphism as 1. In PGLS regression analysis, polymorphism group label is used as the independent variable, and the number of spacer sequences as the dependent variable.

Supplementary Table 4. PGLS Results of the Relationship Between Terminal Repeat Polymorphism and Spacer Number in Short-Repeat (< 38 bp) Genomes

| Polymorphism Threshold | Low Polymorphism | High Polymorphism | Model | Slope | *p* |
| --- | --- | --- | --- | --- | --- |
| 0 | 280 | 523 | Lambda | 2.121  0.989  0.989 | 0.335  0.578  0.578 |
|  |  |  | BM |  |  |
|  |  |  | EB |  |  |
| 1 | 392 | 411 | Lambda | 0.997  1.385  1.385 | 0.632  0.351  0.351 |
|  |  |  | BM |  |  |
|  |  |  | EB |  |  |
| 2 | 476 | 327 | Lambda | 1.232  -0.018  -0.018 | 0.548  0.989  0.989 |
|  |  |  | BM |  |  |
|  |  |  | EB |  |  |
| 3 | 546 | 257 | Lambda | 1.913  0.596  0.596 | 0.359  0.643  0.643 |
|  |  |  | BM |  |  |
|  |  |  | EB |  |  |
| 4 | 620 | 183 | Lambda | 4.901  0.085  0.085 | 0.034  0.955  0.955 |
|  |  |  | BM |  |  |
|  |  |  | EB |  |  |
| 5 | 673 | 130 | Lambda | 5.303  -0.478  -0.478 | 0.045  0.760  0.760 |
|  |  |  | BM |  |  |
|  |  |  | EB |  |  |
| 6 | 720 | 83 | Lambda | 4.342  -0.572  -0.572 | 0.176  0.765  0.765 |
|  |  |  | BM |  |  |
|  |  |  | EB |  |  |
| 7 | 745 | 58 | Lambda | 1.256  -3.389  -3.389 | 0.733  0.105  0.105 |
|  |  |  | BM |  |  |
|  |  |  | EB |  |  |
| 8 | 770 | 33 | Lambda | 0.051 | 0.992 |
|  |  |  | BM | -6.484 | 0.016 |
|  |  |  | EB | -6.484 | 0.016 |

Data includes 803 bacterial genomes.The threshold for polymorphism is set as the number of base differences between the terminal repeat and the consensus sequence, ranging from 0 to 8. Low polymorphism is defined as less than or equal to the threshold, and high polymorphism is defined as greater than the threshold. We labeled groups with low and no polymorphism as 0 and groups with high polymorphism as 1. In PGLS regression analysis, polymorphism group label is used as the independent variable, and the number of spacer sequences as the dependent variable.

Supplementary Table 5. PGLS Results of the Relationship Between Terminal Repeat Polymorphism and Spacer Number in Long-Repeat (≥ 40 bp) Genomes

| Polymorphism Threshold | Low Polymorphism | High Polymorphism | Model | Slope | | | *p* | |
| --- | --- | --- | --- | --- | --- | --- | --- | --- |
| 0 | 3 | 10 | Lambda | 18.212 | | | 0.100 | |
|  |  |  | BM | 16.959 | | | 0.003 | |
|  |  |  | OU | 17.233 | | | 0.022 | |
|  |  |  | EB | | 16.959 | | | 0.003 |
| 2 | 4 | 9 | Lambda | | 21.743 | | | 0.016 |
|  |  |  | BM | | 17.625 | | | 0.002 |
|  |  |  | OU | | 17.638 | | | 0.011 |
|  |  |  | EB | | 17.625 | | | 0.002 |
| 3 | 5 | 8 | Lambda | | 20.635 | | | 0.016 |
|  |  |  | BM | | 17.476 | | | 0.002 |
|  |  |  | OU | | 17.048 | | | 0.012 |
|  |  |  | EB | | 17.476 | | | 0.002 |
| 4 | 6 | 7 | Lambda | | 22.260 | | | 0.003 |
|  |  |  | BM | | 17.997 | | | 5.5×10^-4^ |
|  |  |  | OU | | 18.763 | | | 0.004 |
|  |  |  | EB | | 17.997 | | | 5.5×10^-4^ |
| 6 | 8 | 5 | Lambda | | 18.992 | 0.009 | | |
|  |  |  | BM | | 17.203 | 0.002 | | |
|  |  |  | OU | | 16.125 | 0.017 | | |
|  |  |  | EB | | 17.203 | 0.002 | | |
| 11 | 9 | 4 | Lambda | | 21.835 | 0.004 | | |
|  |  |  | BM | | 17.947 | 7.7×10^-4^ | | |
|  |  |  | OU | | 18.169 | 0.008 | | |
|  |  |  | EB | | 17.947 | 7.7×10^-4^ | | |

Data includes 13 bacterial genomes.The threshold for polymorphism is set as the number of base differences between the terminal repeat and the consensus sequence, ranging from 0, 2, 3, 4, 6, and11. Low polymorphism is defined as less than or equal to the threshold, and high polymorphism is defined as greater than the threshold. We labeled groups with low and no polymorphism as 0 and groups with high polymorphism as 1. In PGLS regression analysis, polymorphism group label is used as the independent variable, and the number of spacer sequences as the dependent variable.

Supplementary Table 6. PGLS Results of the Relationship Between Terminal Repeat Polymorphism and Spacer Number in Short-Repeat (< 40 bp) Genomes

| Polymorphism Threshold | Low Polymorphism | High Polymorphism | Model | Slope | *p* |
| --- | --- | --- | --- | --- | --- |
| 0 | 280 | 524 | Lambda | 2.176 | 0.318 |
|  |  |  | BM | 0.957 | 0.584 |
|  |  |  | OU | 1.228 | 0.558 |
|  |  |  | EB | 0.957 | 0.584 |
| 1 | 394 | 410 | Lambda | 1.141 | 0.582 |
|  |  |  | BM | 1.405 | 0.337 |
|  |  |  | OU | 1.181 | 0.522 |
|  |  |  | EB | 1.405 | 0.337 |
| 2 | 479 | 325 | Lambda | 1.423 | 0.484 |
|  |  |  | BM | -0.005 | 0.997 |
|  |  |  | OU | 0.530 | 0.758 |
|  |  |  | EB | -0.005 | 0.997 |
| 3 | 550 | 254 | Lambda | 1.968 | 0.342 |
|  |  |  | BM | 0.602 | 0.634 |
|  |  |  | OU | 0.875 | 0.606 |
|  |  |  | EB | 0.602 | 0.634 |
| 4 | 624 | 180 | Lambda | 4.961 | 0.031 |
|  |  |  | BM | 0.088 | 0.952 |
|  |  |  | OU | 1.983 | 0.310 |
|  |  |  | EB | 0.088 | 0.952 |
| 5 | 675 | 129 | Lambda | 5.159 | 0.049 |
|  |  |  | BM | -0.484 | 0.753 |
|  |  |  | OU | 2.161 | 0.301 |
|  |  |  | EB | -0.484 | 0.753 |
| 6 | 721 | 83 | Lambda | 4.335 | 0.172 |
|  |  |  | BM | -0.566 | 0.764 |
|  |  |  | OU | 2.646 | 0.299 |
|  |  |  | EB | -0.566 | 0.764 |
| 7 | 746 | 58 | Lambda | 1.277 | 0.726 |
|  |  |  | BM | -3.383 | 0.100 |
|  |  |  | OU | -1.253 | 0.658 |
|  |  |  | EB | -3.383 | 0.100 |

Data includes 804 bacterial genomes.The threshold for polymorphism is set as the number of base differences between the terminal repeat and the consensus sequence, ranging from 0 to 7. Low polymorphism is defined as less than or equal to the threshold, and high polymorphism is defined as greater than the threshold. We labeled groups with low and no polymorphism as 0 and groups with high polymorphism as 1. In PGLS regression analysis, polymorphism group label is used as the independent variable, and the number of spacer sequences as the dependent variable.

Supplementary Table 7. PGLS Results of the Relationship Between Terminal Repeat Polymorphism and Spacer Number in Long-Repeat (≥ 36 bp) Genomes

| Polymorphism Threshold | Low Polymorphism | High Polymorphism | Model | Slope | *p* |
| --- | --- | --- | --- | --- | --- |
| 0 | 142 | 166 | Lambda | 1.679 | 0.538 |
|  |  |  | BM | -0.911 | 0.665 |
|  |  |  | OU | 2.492 | 0.377 |
|  |  |  | EB | -0.911 | 0.665 |
| 1 | 182 | 126 | Lambda | 0.792 | 0.776 |
|  |  |  | BM | 3.436 | 0.118 |
|  |  |  | OU | 3.512 | 0.222 |
|  |  |  | EB | 3.436 | 0.118 |
| 2 | 211 | 97 | Lambda | 2.005 | 0.490 |
|  |  |  | BM | 3.071 | 0.210 |
|  |  |  | OU | 4.832 | 0.120 |
|  |  |  | EB | 3.071 | 0.210 |
| 3 | 229 | 79 | Lambda | 2.350 | 0.446 |
|  |  |  | BM | 3.926 | 0.138 |
|  |  |  | OU | 3.543 | 0.286 |
|  |  |  | EB | 3.926 | 0.138 |
| 4 | 246 | 62 | Lambda | 3.834 | 0.258 |
|  |  |  | BM | 4.222 | 0.132 |
|  |  |  | OU | 5.537 | 0.123 |
|  |  |  | EB | 4.223 | 0.132 |
| 5 | 259 | 49 | Lambda | 3.753 | 0.315 |
|  |  |  | BM | -0.561 | 0.859 |
|  |  |  | OU | 6.126 | 0.127 |
|  |  |  | EB | -0.561 | 0.859 |
| 6 | 272 | 36 | Lambda | 6.564 | 0.123 |
|  |  |  | BM | 9.078 | 0.014 |
|  |  |  | OU | 10.990 | 0.017 |
|  |  |  | EB | 9.078 | 0.014 |
| 7 | 279 | 29 | Lambda | 3.796 | 0.425 |
|  |  |  | BM | 11.035 | 0.012 |
|  |  |  | OU | 7.632 | 0.142 |
|  |  |  | EB | 11.035 | 0.012 |

Data includes 308 bacterial genomes.The threshold for polymorphism is set as the number of base differences between the terminal repeat and the consensus sequence, ranging from 0 to 7. Low polymorphism is defined as less than or equal to the threshold, and high polymorphism is defined as greater than the threshold. We labeled groups with low and no polymorphism as 0 and groups with high polymorphism as 1. In PGLS regression analysis, polymorphism group label is used as the independent variable, and the number of spacer sequences as the dependent variable.

Supplementary Table 8. PGLS Results of the Relationship Between Terminal Repeat Polymorphism and Spacer Number in Short-Repeat (< 36 bp) Genomes

| Polymorphism Threshold | Low Polymorphism | High Polymorphism | Model | Slope | *p* |
| --- | --- | --- | --- | --- | --- |
| 0 | 183 | 415 | Lambda | 2.142 | 0.487 |
|  |  |  | BM | 2.866 | 0.282 |
|  |  |  | OU | 2.073 | 0.504 |
|  |  |  | EB | 2.866 | 0.282 |
| 1 | 266 | 332 | Lambda | 0.911 | 0.744 |
|  |  |  | BM | 0.303 | 0.881 |
|  |  |  | OU | -0.134 | 0.958 |
|  |  |  | EB | 0.303 | 0.881 |
| 2 | 330 | 268 | Lambda | 0.246 | 0.926 |
|  |  |  | BM | -0.892 | 0.595 |
|  |  |  | OU | -0.420 | 0.853 |
|  |  |  | EB | -0.892 | 0.595 |
| 3 | 391 | 207 | Lambda | 2.152 | 0.420 |
|  |  |  | BM | 0.015 | 0.992 |
|  |  |  | OU | 0.539 | 0.806 |
|  |  |  | EB | 0.015 | 0.992 |
| 4 | 451 | 147 | Lambda | 5.037 | 0.087 |
|  |  |  | BM | 0.864 | 0.643 |
|  |  |  | OU | 2.549 | 0.315 |
|  |  |  | EB | 0.864 | 0.643 |
| 5 | 495 | 103 | Lambda | 4.867 | 0.149 |
|  |  |  | BM | 0.874 | 0.645 |
|  |  |  | OU | 2.572 | 0.332 |
|  |  |  | EB | 0.874 | 0.645 |
| 6 | 534 | 64 | Lambda | 4.010 | 0.329 |
|  |  |  | BM | 0.435 | 0.854 |
|  |  |  | OU | 3.022 | 0.358 |
|  |  |  | EB | 0.435 | 0.854 |
| 7 | 558 | 40 | Lambda | 3.612 | 0.460 |
|  |  |  | BM | -1.866 | 0.467 |
|  |  |  | OU | 0.275 | 0.941 |
|  |  |  | EB | -1.866 | 0.467 |

Data includes 598 bacterial genomes.The threshold for polymorphism is set as the number of base differences between the terminal repeat and the consensus sequence, ranging from 0 to 7. Low polymorphism is defined as less than or equal to the threshold, and high polymorphism is defined as greater than the threshold. We labeled groups with low and no polymorphism as 0 and groups with high polymorphism as 1. In PGLS regression analysis, polymorphism group label is used as the independent variable, and the number of spacer sequences as the dependent variable.

Supplementary Table 9. Relationship between CRISPR repeat length and spacer number in bacteria.

| Formula (dependent variable ~ independent variable) | Model | Slope | *p* |
| --- | --- | --- | --- |
| Spacer number ~ Repeat length | BM | -1.178 | 5.0×10^-10^ |
|  | OU | -0.661 | 5.2×10^-5^ |
|  | EB | -1.178 | 5.0×10^-10^ |

Data includes 1,958 bacterial genomes.

Supplementary Table 10. Relationship between spacer number and repeat sequence consistency

| Formula (dependent variable ~ independent variable) | Model | Slope | *p* |
| --- | --- | --- | --- |
| Spacer number ~ Repeat sequence consistency | BM | 0.975 | 1.0×10^-22^ |
|  | OU | 1.411 | 6.6×10^-32^ |
|  | EB | 0.975 | 1.0×10^-22^ |

Data includes 1,958 bacterial genomes.

Supplementary Table 11. Relationship between spacer number and secondary structure stability of consensus repeat sequences

| Formula (dependent variable ~ independent variable) | Model | Slope | *p* |
| --- | --- | --- | --- |
| Spacer number ~ Secondary structure stability of consensus repeat sequences | Lambda | -1.039 | 3.8×10^-4^ |
|  | BM | -1.121 | 5.0×10^-6^ |
|  | EB | -1.121 | 5.0×10^-6^ |

Data includes 814 bacterial genomes.

Supplementary Table 12. Comparing obsolete arrays with functional arrays in spacer number and repeat sequence consistency

| Formula (dependent variable ~ independent variable) | Model | Slope | *p* |
| --- | --- | --- | --- |
| Spacer number ~ Obsolete arrays with functional arrays | BM | 12.457 | 9.7×10^-17^ |
|  | OU | 13.087 | 4.1×10^-11^ |
|  | EB | 12.457 | 9.7×10^-17^ |
| Repeat sequence consistency ~ Obsolete arrays with functional arrays | BM | 3.906 | 3.0×10^-31^ |
|  | OU | 3.439 | 6.4×10^-18^ |
|  | EB | 3.906 | 3.0×10^-31^ |

Data includes 1,920 bacterial genomes.

Supplementary Table 13. Comparing obsolete arrays with functional arrays in Secondary structure stability of consensus repeat sequences

| Formula (dependent variable ~ independent variable) | Model | Slope | *p* |
| --- | --- | --- | --- |
| Secondary structure stability of consensus repeat sequences ~ Obsolete arrays with functional arrays | BM | -0.350 | 0.176 |
|  | OU | -0.464 | 0.140 |
|  | EB | -0.350 | 0.176 |

Data includes 787 bacterial genomes.

Supplementary Table 14. Relationship between spacer number and repeat sequence consistency

| Formula (dependent variable ~ independent variable) | canonical arrays (n = 1753) | | | orphan arrays (n = 167) | | |
| --- | --- | --- | --- | --- | --- | --- |
|  | Model | Slope | *p* | Model | Slope | *p* |
| Spacer number ~ Repeat sequence consistency | BM | 0.903 | **8.7×10^-14^** | Lambda | 0.536 | **0.009** |
|  | OU | 1.433 | **3.0×10^-24^** | BM | 0.169 | 0.236 |
|  | EB | 0.903 | **8.7×10^-14^** | EB | 0.169 | 0.236 |

The results with the *p* value < 0.05 are presented in bold.

Supplementary Table 15. Relationship between spacer number and Secondary structure stability of consensus repeat sequences

| Formula (dependent variable ~ independent variable) | canonical arrays (n = 724) | | | orphan arrays (n = 63) | | |
| --- | --- | --- | --- | --- | --- | --- |
|  | Model | Slope | *p* | Model | Slope | *p* |
| Spacer number ~ Secondary structure stability of consensus repeat sequences | Lambda | -0.904 | **0.004** | BM | 0.022 | 0.956 |
|  | BM | -1.252 | **5.9**×**10^-6^** | OU | 0.418 | 0.245 |
|  | EB | -1.252 | **5.9**×**10^-6^** | EB | 0.044 | 0.910 |

The results with the *p* value < 0.05 are presented in bold.
